# Spatially Multiplexed Single-molecule Translocations through a Nanopore at Controlled Speeds

**DOI:** 10.1101/2022.10.08.511427

**Authors:** S.M. Leitao, V. Navikas, H. Miljkovic, B. Drake, S. Marion, G. Pistoletti Blanchet, K. Chen, S. F. Mayer, U. F. Keyser, A. Kuhn, G. E. Fantner, A. Radenovic

## Abstract

Nanopores are one of the most successful label-free single-molecule techniques with several sensing applications such as biological screening, diagnostics, DNA and protein sequencing^1–4^. In current nanopore technologies, stochastic processes influence both the selection of the translocating molecule, translocation rate and translocation velocity^5,6^. As a result, single-molecule translocations are difficult to control spatially and temporally. Here we present a novel method where we engineer precise spatial and temporal control into the single-molecule experiment. We use a glass nanopore mounted on a 3D nanopositioner to spatially select molecules, deterministically tethered on a glass surface, for controlled translocations. By controlling the distance between the nanopore and the glass surface, we can actively select the region of interest on the molecule and scan it a controlled number of times and at controlled velocity. Decreasing the velocity and averaging thousands of consecutive readings of the same molecule increases the signal-to-noise ratio (SNR) by two orders of magnitude compared to free translocations. We applied our method to various DNA constructs, achieving down to single nucleotide gap resolution. The spatial multiplexing combined with the sub-nanometer resolution could be used in conjunction with micro-array technologies to enable screening of DNA, improving point of care devices, or enabling high-density, addressable DNA data storage.

## Main

Biological nanopores have been used to successfully sequence single-stranded DNA (ssDNA) which is made possible by slowing the speed of the translocation with protein motors^7,8^. However, the stochastic nature of which molecule enters the pore at which time, as well as the residual fluctuation in the feed rate of the protein motor remains challenging^9^.

Solid-state nanopores have been engineered to attempt the same molecular detectability of biological nanopores while keeping full control of the pore design and properties in a stable system^10–13^. Pores can be tuned to the desired size for the molecules in question, enabling the detection and characterization of a broad range of biomolecules such as double-stranded DNA (dsDNA), proteins and DNA-protein complexes^3,14^. However, speed control with molecular motors has not yet been demonstrated with solid-state nanopores. Even though the conductance drop in solid-state nanopores can be higher than in biological nanopores, the uncontrolled translocation speed hampers the potentially higher SNR. Therefore, the free translocation speed limits high fidelity measurements, making single base pair resolution along freely translocating dsDNA impossible to achieve.

Uncontrolled velocity limits the temporal and spatial resolution due to a finite amplifier bandwidth and the noise density integrated over the measurement bandwidth^15^. This limitation decreases the signal-to-noise ratio (SNR), which is critical for detecting small features along the DNA nanostructure^9^. High translocation speeds and low SNR on single readings have this far hampered the use of solid-state devices in the study of complex forms of dsDNA.

Nanocapillaries, referred to as glass nanopores, are well suited for the detection of DNA structures^16,17^ since they have good SNR characteristics for high-bandwidth measurements. Glass nanopores can be manufactured to radii below 10 nm by precisely controlling the diameter of the opening with an electron beam irradiation or by depositing the additional coatings by controllable wet-chemical silanization^18^. They can also be integrated with other techniques, such as optical detection and optical tweezers^19^. The latter allows probing force during translocation but the very high force resolution requires low stiffness of the optical trap and prevents sub-nanometer displacement control.

Scanning Ion Conductance Microscopy (SICM) on the other hand, has very high accuracy displacement control combined with high-bandwidth, low noise current sensing. SICM uses a glass nanopore as a probe to image biological surfaces by moving the nanocapillary with nanometer precision towards a surface while measuring the current to detect the sample topography^20–22^. This microscopy method reveals the dynamics of nanostructures on the cell membrane and can be combined with super-resolution fluorescence techniques^23,24^. Here we utilize the benefits of SICM to overcome the limitations of uncontrolled translocation of DNA through glass nanopores by combining the high SNR and high bandwidth of the glass nanopores with the high degree of displacement control of SICM to achieve accurate, repeated control of single-molecule translocation experiments.

We modified our previously developed HS-SICM setup^23^ with the addition of a closed loop, long range XYZ piezo scanner to accurately control the pipette motion (Figure 1a). Analogously to single-molecule force spectroscopy experiments^25^ performed by Atomic Force Microscopy (AFM), with SICM we can perform current spectroscopy curves on single molecules. We, therefore, refer to our measurements as scanning ion conductance spectroscopy (SICS). In the first step, the pipette is approached within 0.2-1 µm to the surface. The electrophoretic force captures the DNA when the probe approaches the free-end of a tethered molecule (Figure 1b-1). Due to the molecule entering the pore, the conductance through the glass nanopore drops. Then, the glass nanopore is moved up with sub-nanometer precision by the closed-loop piezo (Figure 1b-2). The measured conductance through the nanopore can then be plotted as a function of distance along the molecule (Figure 1b-3), until the molecule is pulled out completely from the nanopore (Figure 1b-4), see conductance plot in red below the method schematics in Figure 1b. After this, we can go to a different molecule at a different XY position or scan the same molecule again. We can even stop before reaching step 4, and measure the same region of the molecule multiple times without removing the molecule from the pore. As a proof-of-concept, we performed SICS mapping of 10 × 10 curves over a 40 × 40 µm² area detecting dCas9 binding specificity along the DNA contour length (SI Figure 1). Using a glass coverslip as a substrate to tether, the single-molecules permits concomitant single-molecule fluorescence measurements in which we labeled dCas9 proteins with Alexa Fluor™ 647 dye (SI Figure 2).

**Figure 1.**
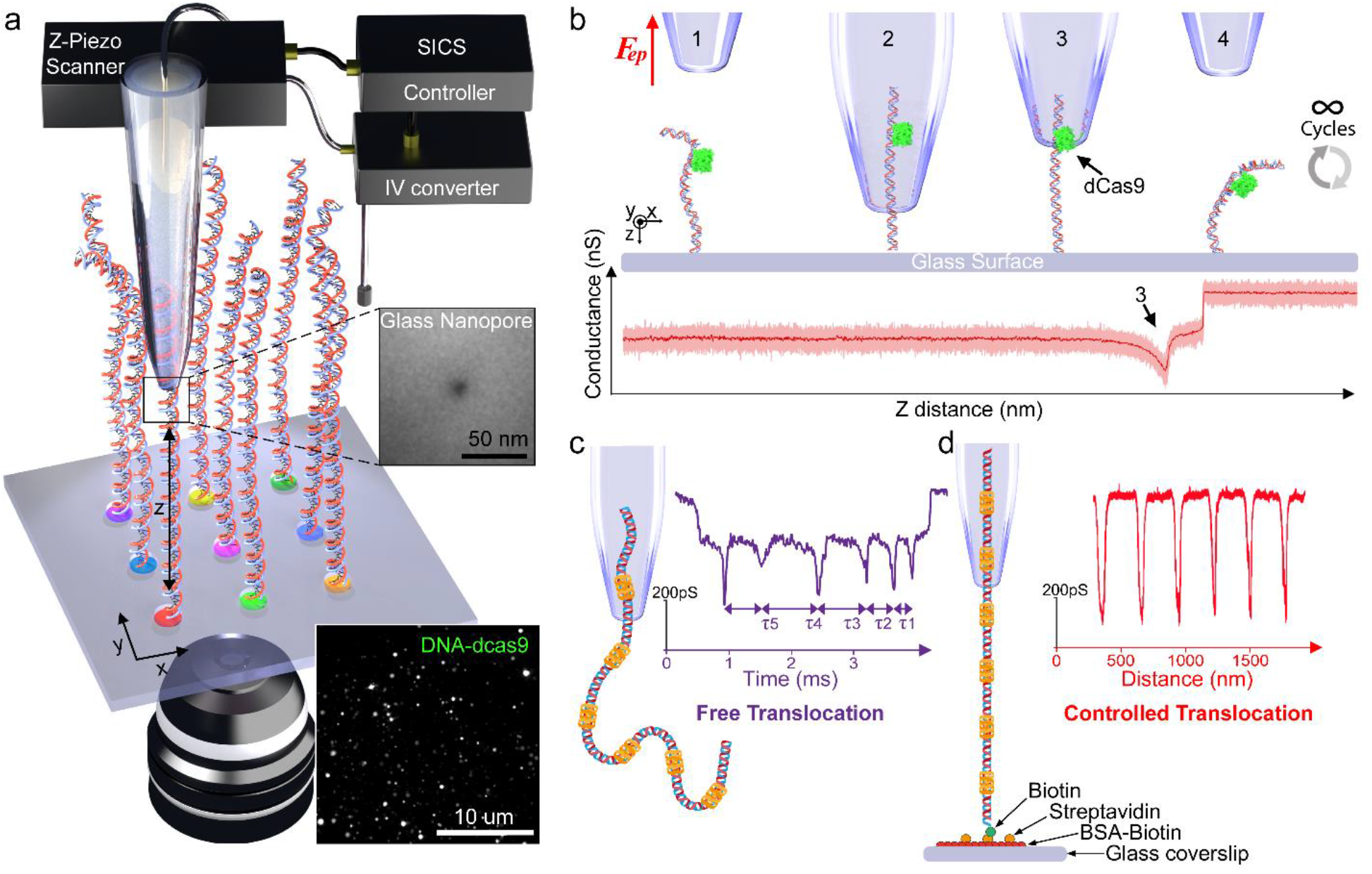
Controlled translocations of single molecules with nanopore-based scanning ion conductance spectroscopy (SICS). **a)** Schematics of the SICS system combined with fluorescence microscopy on spatially multiplexed single molecules (DNA). **b)** Principle of SICS step-by-step. (1): DNA shortly before capture in the nanopore by an electrophoretic force generated by a positive bias (50 – 600 mV). (2): Translocation along the DNA length. (3): Identification of a feature. (4): Complete translocation followed by thousands of potential translocation cycles on the same molecule or mapping out different molecules upon XY displacement. The generated data corresponds to a conductance-distance curve revealing the DNA nanostructure, identifying a DNA-dCas9 complex in this case (red curve). 20 nm glass nanopore radius at 1 µm/s translocation velocity, 200 mV bias in 400 mM KCl, pH=7.4. **c)** Free-translocation signature of a DNA ruler with 8 nm glass nanopore radius in optimal conditions, 600 mV bias in 4 M LiCl, pH=7.4. **d)** SICS controlled translocation signature of a DNA ruler in optimal conditions with 8 nm glass nanopore radius at 1 μm/s translocation velocity, 300 mV bias in 1 M KCl, pH=7.4. DNA molecules are tethered to a glass surface *via* biotin-streptavidin binding.

**Figure 2.**
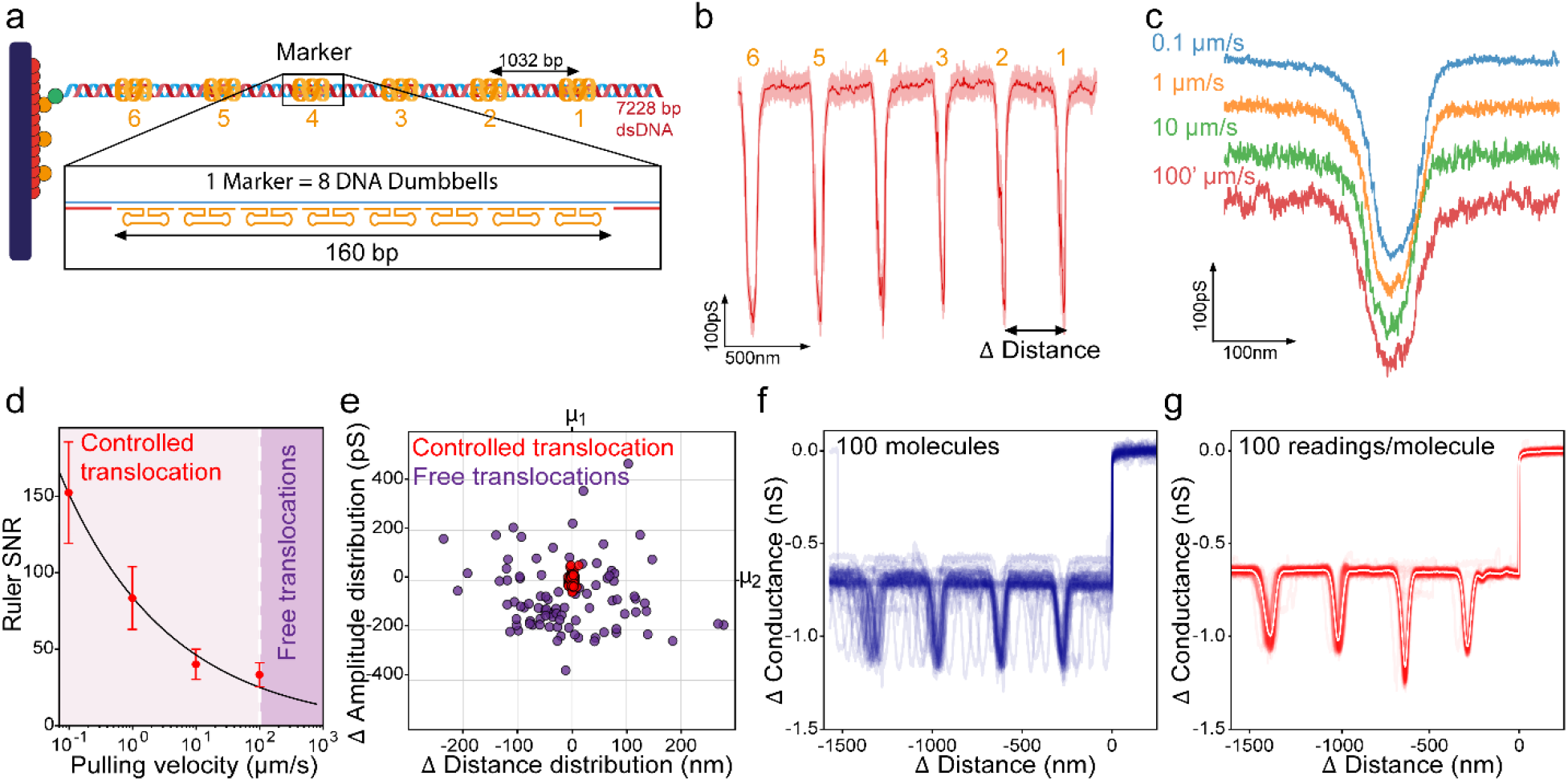
Controlled translocation of custom-designed DNA rulers increases SNR, precision, and accuracy. **a)** Design of the DNA ruler construct, composed of 7’228 base pairs with 6 markers containing DNA dumbbell hairpins separated by equal 1032 bp intervals. **b)** Controlled translocation of DNA ruler with 7 nm radius glass nanopore at 1 µm/s translocation velocity (dark red shows the signal processed by a moving average filter with a 1 nm window size and orange numbers above the trace indicate marker location along the DNA ruler). **c)** Controlled translocation at several velocities: 0.1 μm/s (blue), 1 μm/s (orange), 10 μm/s (green) and 100 μm/s (red); 8 nm radius glass nanopore. **d)** SNR for the detection of marker 1 (First from the free-end) at different translocation velocities (n=10). Error bars correspond to the standard deviation. **e)** Distribution of the amplitude for marker 1 and the distance between markers 1 and 2, in 1 μm/s controlled translocations (red) VS free translocations (purple), with glass nanopores of the same size (8 nm radius), centered to the average distance (µ1) and average amplitude (µ2), n = 100. **f)** Overlay of controlled translocations for 100 different molecules tethered on the surface at 1 μm/s translocation velocity. Molecules were aligned to the DNA exit. **g)** Overlay of controlled translocation of the same molecule (red) and their average (in white), n=100; 8 nm radius glass nanopore at 1 μm/s translocation velocity. All controlled translocation experiments were performed with 300 mV bias in 1 M KCl, pH=7.4.

The key difference between our technique and traditional nanopore experiments is how the translocation speed is controlled. In conventional nanopore experiments, the speed of the translocation is determined by the electrophoretic force and the viscous drag of the molecules in the solution and the pore. The speed of translocation depends on many factors, such as ionic strength, molecular charge, bias voltage, and nanopore geometry. The result is a stochastic nature of the translocation. This manifests itself in non-uniform dwell-time distribution between the detection of equally spaced motifs on a molecule such as a DNA ruler, see Figure 1c (free-translocation) and SI Figure 3 and 4. Free translocations generate conductance signals as a function of time with non-uniform translocation velocity^17,26^. In addition to stochastic effects influencing the translocation speed, also the selection of which molecules are translocated is governed by stochastic factors.

**Figure 3.**
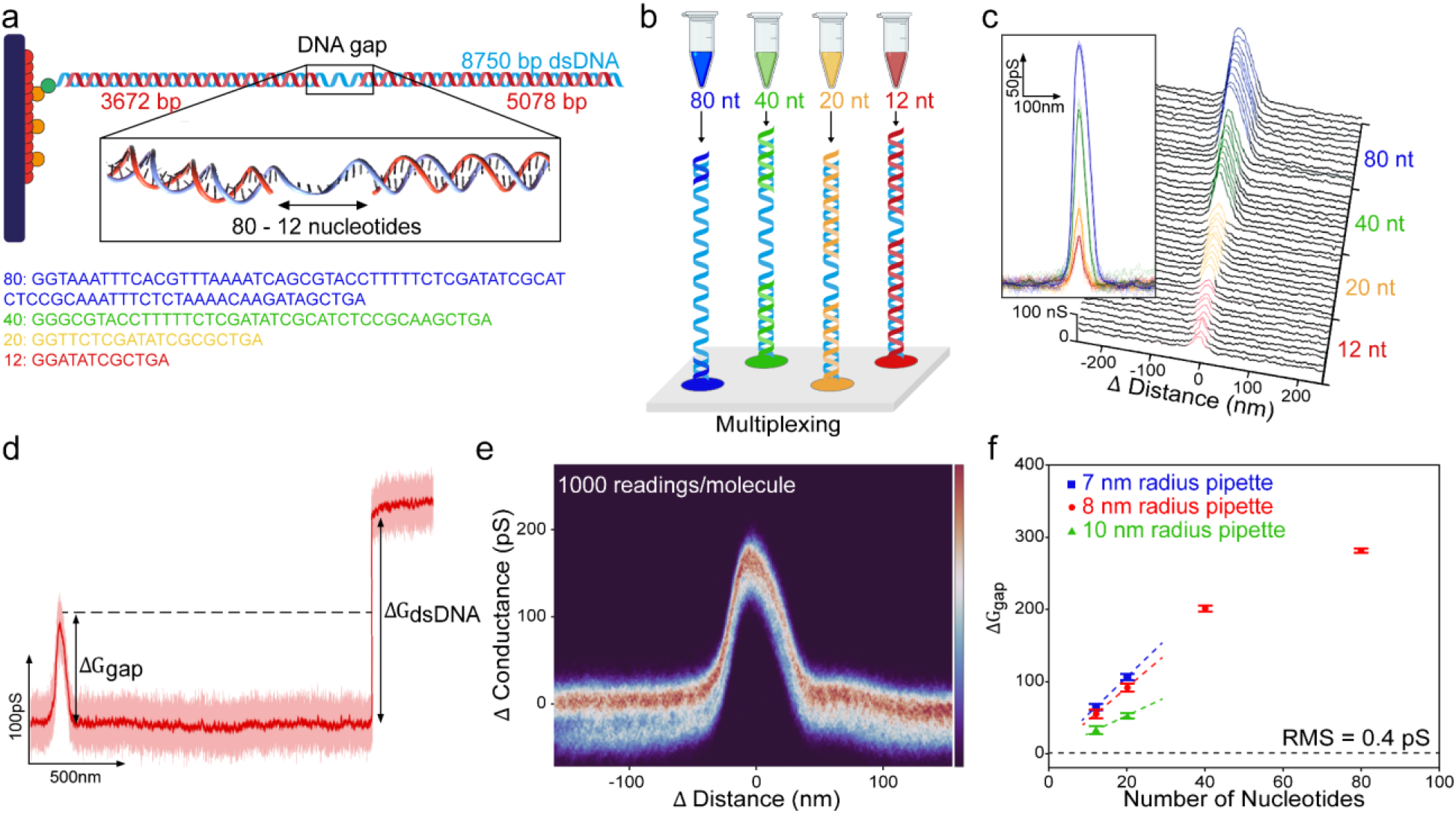
Spatial addressability of single-molecule translocations demonstrated on DNA gaps. **a)** Design of DNA gap construct composed of 8’750 base pairs and containing a central single-stranded region (“gap”) of 4 different sizes: 80, 40, 20, or 12 nucleotides. **b)** Spatially multiplexed DNA gap molecules. **c)** Controlled translocation signals of DNA constructs with the different nucleotide gaps (n=10 on the same molecule with the average in dark color) and corresponding waterfall plot. 8 nm glass nanopore radius at 1 μm/s translocation velocity. **d)** Controlled translocation of the construct with an 80 nucleotide gap. The conductance amplitude (ΔGgap) of half the dsDNA translocation amplitude (ΔGdsDNA); 12 nm glass nanopore radius at 1 μm/s translocation velocity (Dark red). **e)** Probability density map of 1000 readings from the same 80 nucleotide gap, acquired at 10 nm glass nanopore radius at 1 μm/s translocation velocity (4 readings/s); the colormap represents the normalized probability of occurrence for a conductance value at a corresponding distance. **f)** Amplitude of the detected gaps (ΔGgap) vs gap length (in nucleotides) with different pipettes. Error bars correspond to the standard deviation (n=10). All the experiments were performed with 300 mV bias in 1 M KCl, pH=7.4.

**Figure 4.**
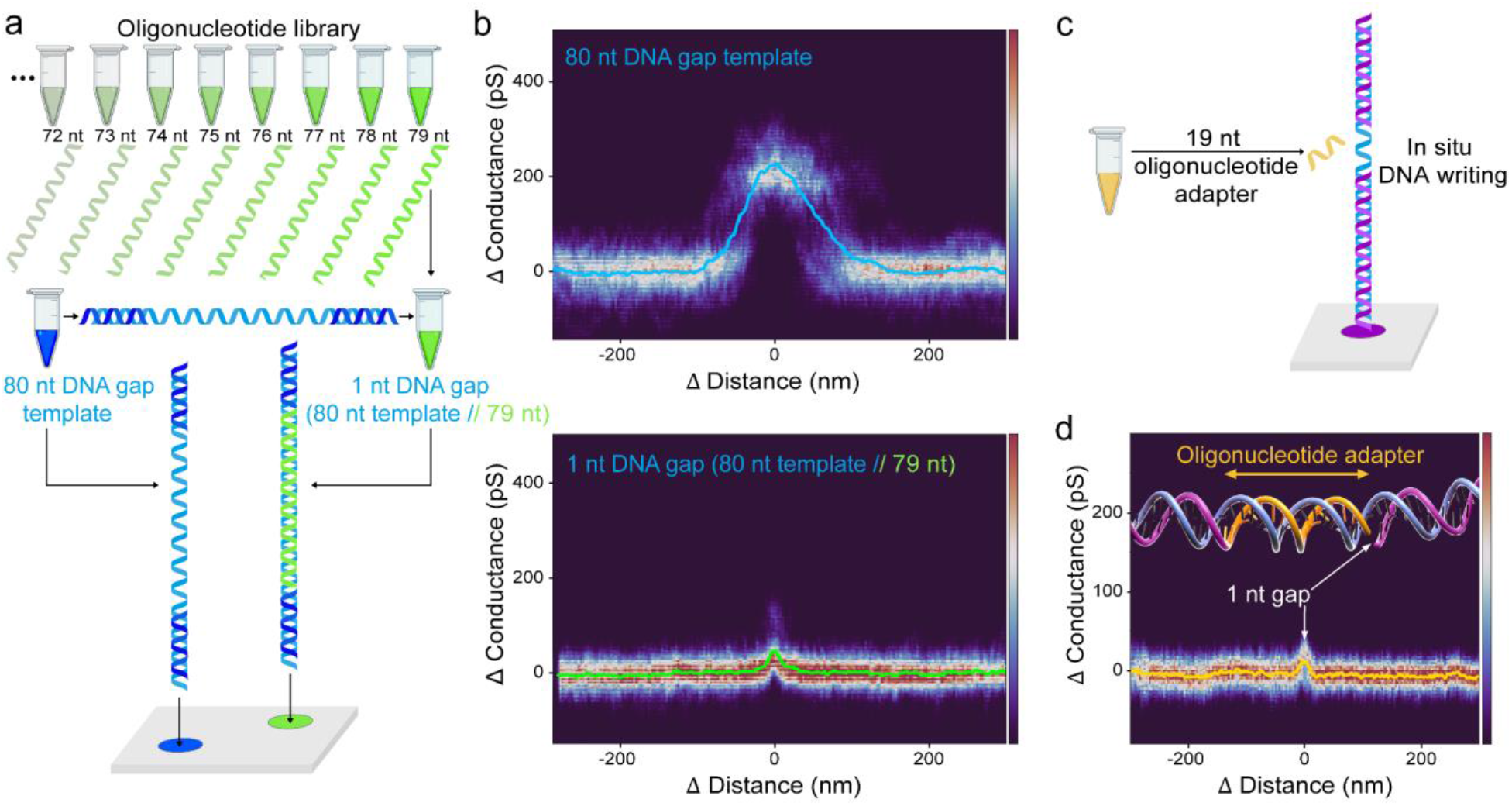
Detection of single nucleotide gaps with SICS. **a)** Strategy for creating shorter, variable gap sizes using multi nt gap template and complementary oligonucleotide library. **b)** Detection of an 80 nt DNA gap template (top) and a single nucleotide gap from a complementary 79 nt-long oligonucleotide hybridized to the gap (bottom), with a 7 nm glass nanopore radius at 1 μm/s translocation velocity. The colormap represents the normalized probability (n=30). **c)** In situ hybridization concept for DNA writing, showing a gap template and addition of the complementary oligonucleotide (gap adaptor) **d)** Reading of a 1 nt gap with 8 nm glass nanopore radius at 1 μm/s translocation velocity. A complementary 19 nt oligonucleotide was hybridized in situ on a 20 nt gap. The colormap represents the normalized probability (n=10). All the experiments were performed with 300 mV bias in 1 M KCl, pH=7.4.

In SICS, however, the molecule of interest is tethered to the surface and the electrophoretic force is countered by a reaction force on the tether, which prevents the molecule from fully translocating. Translocation is solely governed by the motion of the glass nanopore, which can be controlled accurately during the experiment. With the ability to deterministically control the velocity, the conductance traces are now a function of distance instead of time. Figure 1c (controlled translocation) shows the equidistant motif on the DNA ruler. This decouples important experimental parameters (ionic strength, bias voltage) as well as uncontrollable factors (such as nanopore geometry) from the translocation speed and detection bandwidth. We can, therefore, independently optimize experimental parameters to improve the detection limit.

In order to assess the benefits of SICS we used DNA rulers^26,27^ to characterize the effect of translocation velocity on SNR, precision and repeatability. This DNA ruler is composed of a 7’228 base pair DNA strand (2’458 nm) and 6 markers of DNA dumbbell hairpins which are positioned at 1’032 base pairs intervals along the DNA contour (Figure 2a). Figure 2b shows the conductance-distance curve generated by SICS, revealing 6 markers. Controlled translocation with SICS allows decreasing the speed by more than 4 orders of magnitude compared with typical free translocations, leading to an increased SNR (Figure 2c). At the lowest velocity of 0.1 μm/s, the SNR measured was 152 ± 33 with an 8 nm glass nanopore radius (Figure 2d). This corresponds to an order of magnitude improvement of SNR compared with a free-translocation (>100 μm/s velocity). Figure 2e shows a location precision of 1% with controlled translocations, compared with 30% in free translocations with glass nanopores of the same pore size. SICS inherently has lateral control and can map out diverse molecules tethered on the surface or scan the same molecule again. Figure 2f shows the overlay of 100 distinct molecules of the DNA ruler and Figure 2g shows 100 readings of the same selected molecule. This spatial addressability allows us to multiplex the measurements to multiple molecular species.

To take advantage of our SICS platform in terms of spatial addressability, multiplexing and resolution assessment, we engineered dsDNA that contained single-stranded regions (“DNA gaps”) of various sizes. We custom-designed four different DNA constructs (SI Figure 5) that were 8’750 base pairs in length (2’975 nm) and presented DNA gaps of 80, 40, 20, or 12 nucleotides, corresponding to lengths of 27.2, 13.6, 6.8, and 4.1 nm, respectively (Figure 3a). As a proof-of-principle for spatially multiplexed single molecules, we spotted the four different DNA gap constructs in a 2×2 array (Figure 3b). Within a single experiment, using one and the same glass nanopore, we detected and discriminated the four different DNA molecules. Figure 3c shows distinct conductance amplitudes (ΔGgap) for the different 80, 40, 20, and 12 nucleotides (nt) gaps. Figure 3d shows that the measured conductance amplitude of the 80 nt gap (ΔGgap) was half of the dsDNA conductance drop (ΔGdsDNA), as expected. For smaller gap sizes, the measured ΔGgap decreases due to convolution effects. To obtain reliable measurements on the smaller gaps, multiple measurements are desirable.

We use the sub-molecular spatial addressability (position the glass nanopore along the single molecule) to scan the same molecular region multiple times, with tunable bias and bi-directional readings (SI Figure 6 and 7). Figure 3e shows the probability density map of 1000 readings on the same feature (80 nt DNA gap). By recording and averaging conductance-distance curves of the same feature we decreased the RMS noise of the averaged signal from 11.2 pS to 0.4 pS, increasing the SNR 20 times compared with a single reading (SI Figure 8). Moreover, multiple reads of the same feature decreased the error of the measurement, and obtained an average amplitude of 157.3 pS with a standard error of 0.05 pS (0.03 % error), see SI Figure 8. Thus, our method yielded an SNR of 394 on the DNA gap, which corresponds to an improvement of two orders of magnitude compared to free translocations. By accessing specific regions of the DNA molecule and by averaging the conductance fingerprints, we reliably detected gaps as small as 12 nucleotides (4.1 nm) with high SNR for several pipette radii below 10 nm, hinting at an ultimate resolution capability down to single nucleotide gaps (Figure 3f).

To have finer control over the gap size, we created a library of oligonucleotides complementary to the DNA gaps (gap adaptors shown in SI Table 1). As a proof-of-concept, we hybridized 79 nt single-stranded DNA (oligonucleotide) to complementary 80 nt DNA gap molecules (DNA gap template) and then spotted next to the non-hybridized 80 nt DNA gap template (Figure 4a). Within a single experiment and the same glass nanopore, we compared controlled translocations on the 80 nt gap template (Figure 4b - Top) and the 1 nt gap in the hybridized complex (Figure 4b - Bottom). Our results clearly demonstrate the single nt gap resolution of SICS, with an average gap amplitude detected of 31.2 pS for a glass nanopore of 7 nm radius. Furthermore, the hybridization does not have to happen before spotting but rather in situ after spotting the template (Figure 4c). We demonstrated in-situ hybridization of 19 nt oligo strand with a complementary 20 nt gap, and the subsequent successful detection of the single nucleotide gap with an average gap amplitude detected of 16.9 pS for a glass nanopore of 8 nm radius (Figure 4d). This indicates potential applications in diagnostics, DNA data storage and data retrieval^28^.

## Discussion

We achieved full control of the translocation speed providing precise spatial and temporal control of the single-molecule experiments. This allows reading the same molecule, or region of a molecule, thousands of times, as well as scanning an array of different types of molecules. This control is independent of experimental parameters (ionic strength, bias voltage) as well as uncontrollable factors (such as nanopore geometry) and it allows for independent optimization of experimental parameters to improve the detection limit, unlocking the use of nanopore sensing in applications that previously required sub-nanometer resolution. For the first time, it was possible to reproducibly scan selected areas of a molecule, thereby addressing specific molecular regions of interest with unprecedented resolution and throughput. We achieved 100’000 readings per experiment and a scanning rate of 4 readings/s. The ability to perform experiments with different molecular species within one experiment (see Figure 3b), drastically increases the experimental throughput and permits accurate comparison of the results given the identical experimental conditions (same glass nanopore, velocities, and buffer conditions). Besides enabling more detailed biophysical studies, this addressability enables new conceptual approaches to multiplexed diagnostics (multiple analytes on one DNA template), or DNA data storage, where the sample area could be spatially divided into sectors and folders, with the data stored in individual molecules acting as files.

Currently, we used glass capillaries shrunken down with SEM to nanopore size. Fabricating these glass nanopores in a reproducible way remains a challenging task. It offers, however, the possibility to tune the nanopore size depending on the application. Whereas we preferred smaller nanopores for our DNA template-based (DNA rulers, DNA gaps) experiments, DNA/protein complexes often require bigger pores. The conical shape of the glass nanopores intrinsically limits the axial resolution of our measurements. Nevertheless, through the drastic increase of SNR we were able to detect down to single nucleotide gaps in dsDNA. Additionally, SICS measurements could benefit from the fact that our SICS setup is integrated with a state-of-the-art optical microscope, enabling Förster resonance energy transfer (FRET)^29^, single-molecule localization microscopy (SMLM)^30^ and DNA-PAINT (DNA-based Point Accumulation for Imaging in Nanoscale Topography)^31^ experiments.

Compared to other nanopore detection techniques, glass nanopores are more difficult to parallelize. This drawback, however, is partially mitigated by the ability to do spatial multiplexing, meaning that one glass nanopore could probe thousands of molecules arranged in a microarray. The basis of SICS, being the mechanical control of the translocation, could potentially also be realized using solid state nanopores in MEMS devices.

Integration of biological nanopores into the opening of the glass capillary^32^ could combine the benefits of biological nanopores (high resolution and reproducibility) with the benefits of SICS (molecule independent speed control and multiple readings per molecule). We expect this to greatly enhance the suitability of nanopores for the sequencing of peptides and proteins.

## Methods

### Experimental setup

#### SICS system and controlled translocation measurements

The SICS system is based on a high-speed scanning ion conductance microscope (HS-SICM) that we developed recently^1^. We modified our HS-SICM with the addition of a closed loop, long-range XYZ piezo scanner to precisely control the pipette motion and perform mapping curves in a similar way as done in AFM^2^. This enabled us to spatially select single molecules to translocate them at a controlled speed. In order to capture selected molecules, a bias is applied between two Ag/AgCl electrodes across the glass nanopore used in our system. When a positive bias is applied on the negatively charged DNA molecule tethered on the glass surface, the resulting force will pull the molecule through the nanopore, until the molecule is stretched between the surface and the nanopore. By moving the nanopore with respect to the surface, the conductance signature of the molecule inside the nanopore can be measured as a function of distance along the molecule, detecting specific features on the single molecule. The 3D nanopositioner used to translocate the molecules through the nanopore is a piezo-nanopositioning stage with 10 µm Z travel range and 100 µm X-Y travel range (P-733 Piezo NanoPositioner, Physik Instrumente), driven by a low-voltage piezo amplifier (E-500 Piezo Controller System, Physik Instrumente). The 3D nanopositioner was assembled in a custom-built micro translation stage, mounted atop an inverted Olympus IX71 microscope body. Thus, the SICS system enabled correlative fluorescence microscopy with a four-color (405 nm, 488 nm 561 nm, 647 nm) pigtailed Monolithic Laser Combiner (400B, Agilent Technologies), controlled by a custom-written LabVIEW software. An integrated sCMOS camera (Photometrics, Prime 95B) and Micromanager software were used to acquire images. The controlled translocation current signal was amplified by a transimpedance amplifier NF-SA-605F2 (100 MΩ gain, NF corporation) with a bandwidth of 10 kHz. The SICS controller was implemented in LabVIEW on a NI USB-7856R OEM R Series (National Instruments, Austin, TX, USA), to perform high-precision conductance-distance measurements and spatial mapping (conductance-volume mapping), similar to AFM force volume mapping^3^. The controlled translocation signal was acquired at 1 MHz sampling rate. All the controlled translocations shown in this paper were generated in 0.2 - 1 M KCl solution for bias ranging from 50 mV to 600 mV.

#### Data processing and analysis

Controlled translocation curves were processed with a custom-written Python program which was used to automatically filter data, align different molecules, and calculate signal parameters. Single conductance-distance curves were averaged by a moving average filter with a 1 nm window size (Single readings are shown in Figure 1b, 2b and 3d in dark red). The root-mean-square conductance noise (RMS) measured corresponds to the square root of the average squared value of the conductance fluctuations from the mean conductance in a 20 nm range. To calculate the precision and compare controlled translocations with free translocations we used the relative standard deviation: RSD = (standard deviation ÷ average) × 100. Fluorescence image data analysis was performed using Fiji software^4^ and AFM images were processed with Gwyddion software^5^.

### Probe preparation

#### Fabrication and characterization of nanocapillaries

Nanocapillaries were fabricated using a CO2-laser puller (P-2000, Sutter Instrument). Quartz capillaries with 0.5 mm outer diameter and 0.2 mm inner diameter were bought from (Hilgenberg GmbH). Before the pulling process, all capillaries were cleaned with 99 % acetone, 99 % ethanol, MiliQ water (Millipore Corp) and again with 99 % ethanol by sonication in each solution for at least 10 min. After washing, nanocapillaries were dried in a desiccator for 1-2 h until they were completely dry and cleaned for 10 min in oxygen plasma. After the fabrication, nanocapillaries were characterized by a scanning electron microscope (Zeiss, Merlin). Nanocapillary diameters were confirmed using SEM, and as expected, under relatively high imaging current (400 pA), capillaries under 40 nm shrunk due to electron beam heating induced effects^6^ Diameters of all nanocapillaries were measured manually based on SEM images, using Fiji software^4^.

#### Nanocapillary filling procedure

After SEM imaging, nanocapillaries were placed on the cover-glass with double-sided polyimide (Kapton) tape or a specially designed holder fabricated from PEEK plastic. Nanocapillaries were then cleaned with oxygen plasma (Femto A, Diener electronic GmbH) for 300 s at the maximum power setting. Immediately after, the nanocapillaries were immersed in a 400 mM or 1000 mM KCl solution and placed inside the desiccator connected to a vacuum pump. Nanocapillaries were kept under low-pressure (1-10 mbar) for 10 min to pre-fill them and avoid the formation of air bubbles in the thick end of the capillaries. Then they were imaged with an inverted brightfield microscope to confirm pre-filling. After, the nanocapillaries were placed inside a microwave oven (MW 1766 EASY WAVE, P = 700 W, λ= 12.23 cm). The highest power setting was always used. Microwave radiation was applied in heating cycles to heat the solution until its boiling point. The heating duration varied based on the volume and temperature of the solution. The first heating phase took (30-60 s), and subsequent heating phases were significantly shorter (5-10 s) due to the increased temperature of the capillary immersion solution. Heating was always stopped at the boiling point of a solution in order to minimize evaporation^6^. Short 10-20 s pauses were made in between heating steps to allow for the gas to dissolve into the solution. At least 3 heating cycles were performed to complete the filling of the nanocapillary batch^7^. After the procedure, capillaries were kept at 4 °C and the buffer was exchanged after a few hours to ensure the salt concentration was not affected by evaporation. For storage, nanocapillaries were kept at 4 °C in sealed chambers for up to one year.

### Sample preparation

#### Lambda DNA preparation

10 kilobase long λ-DNA was prepared from full-length phage λ-DNA (New England BioLabs) by performing polymerase chain reaction (PCR) using one primer (Microsynth) with a biotin tag on the 5’ end and the second one without biotin at the 3’. PCR was performed using a LongAmp DNA polymerase (New England BioLabs) following the protocol from the manufacturer. The reaction mixture was purified using PCR and a Gel Cleanup kit (Qiagen) from the agarose gel according to the protocol from the manufacturer. The length of the 10 kb λ-DNA product was verified with an agarose gel electrophoresis and the concentration was measured with a NanoDrop 1000 spectrometer.

#### gRNA preparation and immobilization

10kb long λ-DNA was screened for the presence of PAM motifs 5′X20NGG3′ and two targets separated by 5374 bp were selected. Single guide RNAs (gRNAs) were designed to be complementary to the 2 adjacent 20 bp PAM motifs previously selected on the λ-DNA. gRNAs were prepared by in vitro transcription of dsDNA templates carrying a T7 promoter sequence. Transcription templates were generated by PCR amplification of ssDNA templates containing the T7 binding site, 20 bp sequence complementary to the DNA target site, and sgRNA scaffold sequence using Phusion High-Fidelity DNA Polymerase (New England Biolabs). The gRNAs were synthesized by *in vitro* transcription using the MEGAshortscript T7 Transcription kit (Thermo Fisher) according to the manufacturer’s conditions. The gRNAs were treated with Turbo DNase (Thermo Fisher) and MEGAclear Transcription Clean-Up Kit (Thermo Fisher) was used for purification according to the protocol provided by the manufacturer. The concentration of purified gRNAs was measured with a NanoDrop 1000 spectrometer and the length and quality of the sgRNA were estimated by agarose gel electrophoresis. sgRNA samples were kept at -20 °C for a maximum time of 48h until further use.

#### dCas9-DNA complex formation

A commercially available inactive mutant of Cas9 nuclease (dCas9) with N-terminal SNAP-tag (EnGen Spy dCas9) was acquired from New England Biolabs. The protein was labeled with SNAP-Surface Alexa Fluor 647 labeling kit (New England Biolabs) according to the protocol provided by the manufacturer. The unreacted substrate was removed by the size exclusion column. Finally, fluorescently labeled dCas9 was incubated with gRNA in 1× NEBuffer 3.1 (New England Biolabs). 1 nM dCas9 was incubated with 10 nM gRNA for 30 min at 37 °C on an orbital shaker. 0.5 units of RNase inhibitors (Thermo Fisher) were used to prevent the degradation of RNA. After incubation, DNA was added at the final concentration of 50 pM and the mixture was further kept for 30 min at 37 °C on an orbital shaker. Schematics of the final constructs are shown in SI Figure 1 and dCas9-DNA binding controls were performed with AFM (SI Figure 9).

#### DNA ruler construct

DNA ruler constructs were prepared in the group of Prof. Ulrich F. Keyser, using a well-established protocol^8^. The DNA ruler was synthesized by cutting circular 7249 base M13mp18 ssDNA (New England Biolabs) using the enzymes EcoRI and BamHI to form a linear ssDNA chain 7228 bases in length. The final scaffold was then purified and mixed in a 1:5 ratio with 212 oligonucleotides that formed a double-strand with six equidistant zones of dumbbell hairpins. The mixture is then annealed and purified. The final construct contains 6 markers with 8 dumbbells each. Markers are equally positioned in intervals of 1032 bp (See SI Figure 3).

#### DNA gap construct

DNA gap constructs were prepared in the group of Prof. Dr. Alexandre Kuhn. DNA gap constructs were generated using modified 9 kb plasmids. In short, the pPIC9K plasmid (Invitrogen) was mutated using Q5 Site-Directed Mutagenesis (New England BioLabs) to insert two restriction sites specific for the nicking endonuclease Nt.BbvCI (New England BioLabs). Four plasmids were engineered with restriction cut sites that were 80, 40, 20, or 12 nt apart and that flanked an EcoRV restriction site. To generate a specific DNA gap fragment, the corresponding purified plasmid was first linearized by restriction digestion and biotin-labeled using Biotin-11-dUTP (ThermoFisher) and Klenow fragment (New England BioLabs). The DNA fragments were then digested with Nt.BbvCI to obtain single-strand breaks (nicks) and gapped by repeated (3x) heating cycles (90 °C, 60 sec; 60 °C, 10 min; 37 °C, 20 min) in the presence of a single-stranded competitor oligonucleotide (i.e. complementary to the DNA sequence comprised between the nicks)^9^. Finally, the construct was digested with EcoRV to eliminate ungapped fragments^10^ analyzed on a 1 % agarose gel and purified (Monarch DNA gel extraction kit, New England BioLabs). The final dsDNA constructs were 8’750 base pairs long with biotin attached on one end, and a gap of 80, 40, 20, or 12 nucleotides (**See Figure 3** and SI Figure 5). The sequence of the single-stranded stretches in the gap constructs were: 80 nt: GGTAAATTTCACGTTTAAAATCAGCGTACCTTTTTCTCGATATCGCATCTCCGCAAATTTCTCTAAAACAAGATAGCTGA; 40 nt: GGGCGTACCTTTTTCTCGATATCGCATCTCCGCAAGCTGA; 20 nt: GGTTCTCGATATCGCGCTGA; 12 nt: GGATATCGCTGA. Occasionally we observed hairpins during SICS experiments in samples measured right after incubation at 4**°**C (See SI Figure 10).

#### Oligonucleotide-DNA gap template construct

In order to create gaps of shorter length, we created a library of complementary DNA oligos of different lengths that were hybridized on already existing dsDNA with a gap (See SI Table 1). In this way, we got gaps of smaller sizes that can be found in the table. We hybridized them by mixing the ∼9 kb dsDNA gap templates (20, 40, 80) with the oligo (termed gap adaptor) at a 1:100 ratio, and placing it in a thermocycler with the following program: the sample was heated at 70 °C for 5 min and afterward ramped down 1 °C for 1 min per cycle, over 60 cycles.

#### Imaging chambers for DNA immobilization

The imaging chambers that are compatible with single-molecule fluorescence and single-molecule scanning ion conductance microscopy imaging were fabricated from high precision No. 1.5 borosilicate 25 mm coverslips (Marienfeld). Coverslips were cleaned with ethanol, then MiliQ water, dried with nitrogen flow, and cleaned with oxygen plasma (Femto A, Diener electronic GmbH) at maximum power for 660 s. After cleaning the open circular chamber made from a 1 mL pipette tip was glued on top with polydimethylsiloxane (PDMS), forming the final assembly. Fabricated chambers were kept in sealed petri-dishes until further use. For multiplexing, we made four small chambers (each of 5 μl volume) located on the same coverslip (see Figure 3). In the same way, adopting the robotic spotting methodology used in the preparation of microarrays^11^, on a single 25 mm coverslip, one could accommodate more than 400 000 samples in the same buffer conditions.

#### Surface immobilization

100 µL of 1 mg/ml of BSA-Bt (Sigma-Aldrich) in PBS was incubated for at least 1 h in plasma-cleaned chambers to achieve the full glass surface coverage. Samples then were washed 10× with PBS by exchanging half of the solution, but without drying a surface. Samples then were incubated with 0.1 mg/ml of streptavidin (Sigma-Aldrich) for 1 h, followed by a 10× wash with PBS. Finally, 10-100 pM of DNA molecules were incubated for 1 h, followed by 10× wash with PBS or 3.1 NEBuffer. Samples were kept at 4 °C until use. The solution in the imaging chambers was exchanged by performing an additional 10× wash before imaging. The final imaging solutions are described in the imaging and SICS sections. We noticed that samples have not shown signs of degradation even after one year if kept at 4 °C in sealed petri-dishes that eliminates water evaporation.

#### Single-molecule fluorescence imaging

To optimize the surface density of DNA and to perform a control of dCas9 binding, samples were first imaged with a single molecule fluorescence microscope^12^ (See SI Figure 11). For single-molecule imaging and to prevent photobleaching, a reductive/oxidative system (ROXS) based on glucose oxidase and catalase was used. Final buffer composition consisted of 2 mM TROLOX, 40 mM TRIS, 400 mM KCl, 1 % of glucose, 120 units/ml of glucose catalase, 15 units/ml. Buffer was filtered with a 200 nm filter. Imaging was performed in a sealed chamber to avoid oxygen exposure. All chemicals were acquired from (Sigma-Aldrich) unless stated otherwise.

## Data and code availability statements

The codes and data that support the plots in this paper and other findings of this study are available from the corresponding authors upon reasonable request

## Acknowledgements

S.M.L. and G.E.F. acknowledge the support from Swiss Commission for Technology and Innovation under the grant CTI-18330.1., and the European Research Council under grant number ERC-2017-CoG; InCell. V.N., H.M., and A.R. acknowledge the support from National Center of Competence in Research (NCCR) Bio-Inspired Materials and Max-Planck-EPFL Center of Molecular Nanoscience and Technology. K.C. and U.F.K. were funded by PoreDetect (ERC-2019-POC, 899538) and EarlyPore (ERC-2022-POC1, 101069324).

## Author contributions

S.M.L. developed the SICS system and performed the SICS measurements. S.M.L. and V.N. performed data analysis with S.M. support. V.N. and H.M. prepared the single-molecule samples and fabricated nanocapillaries. G.P. and A.K. created DNA gap molecules. H.M. and S.F.M. created the oligonucleotides/DNA gap templates. K.C. and U.K. created DNA rulers and provided free translocation data. S.M.L. and B.D. build the original SICM setup used in this work. A.R., G.E.F., S.M.L. and V.N. designed experiments with input from all the authors. A.R. conceived the method. G.E.F. and A.R. supervised the project. S.M.L., G.E.F., and A.R. wrote the manuscript with input from all the authors.

## Supporting Figures

**Supplementary Figure 1.**
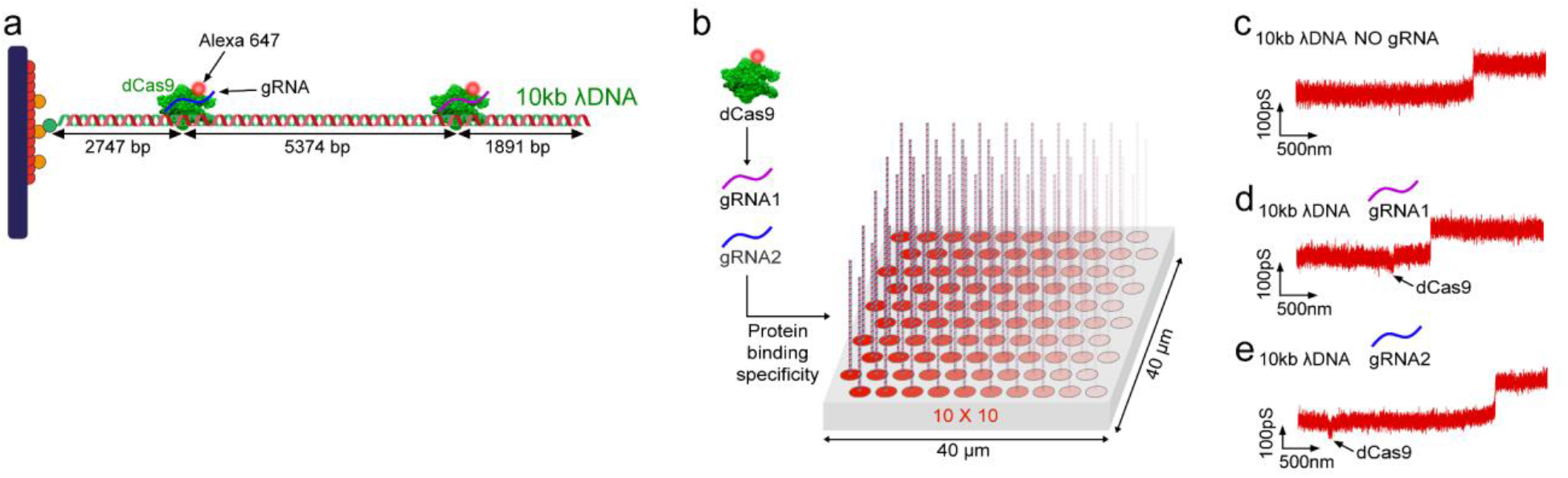
Binding specificity of dCas9 on DNA with SICS. **a)** Design and construct of 10 kb long λ-DNA to test dCas9-DNA complexes with different RNA guides (gRNA). **b)** SICS mapping (10 × 10) of DNA-dcas9 complexes over a 40 × 40 μm^2^ area detecting dCas9 binding specificity. In this proof-of-principle measurement, two gRNA were tested. **c)** SICS conductance-distance curve of a λ-DNA molecule translocated without dCas9/gRNA. **d)** Shows the detection of dCas9/gRNA1. **e)** Shows the detection of dCas9/gRNA2. These experiments were performed with 200 mV bias in 0.4 M KCl, pH=7.4.

**Supplementary Figure 2.**
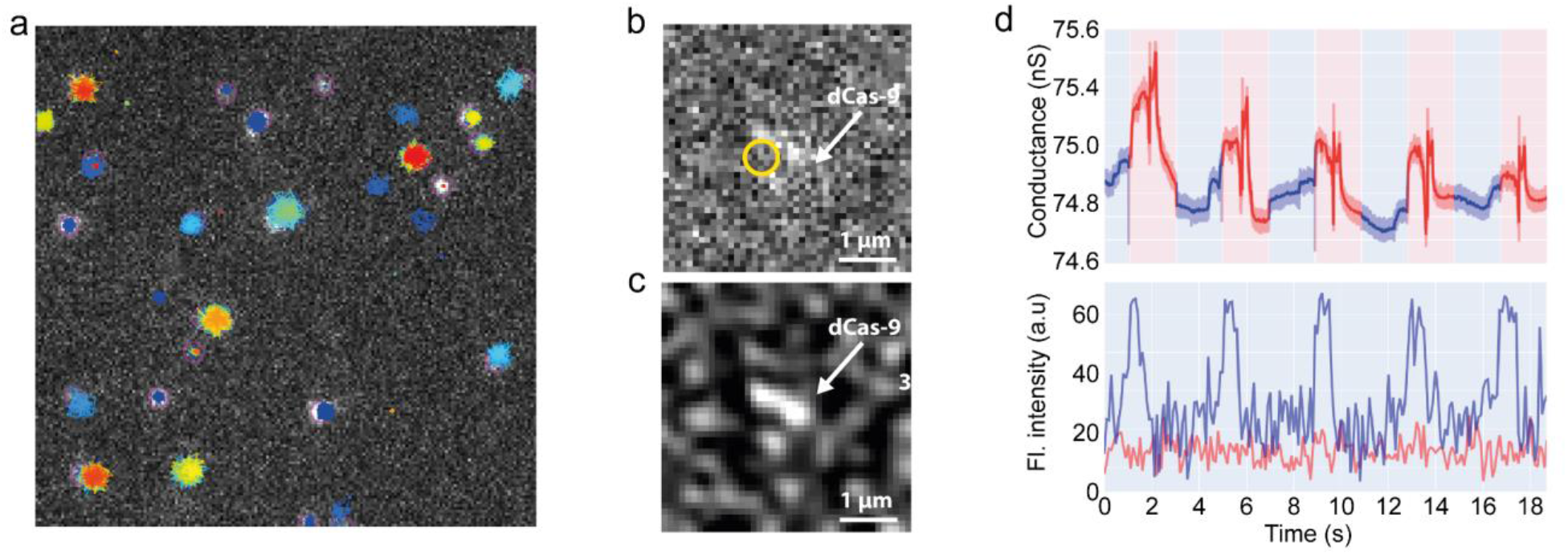
Correlative single-molecule fluorescence and SICS measurement. A single widefield image showing the surface-immobilized DNA-dCas9 complex labeled with Alexa-647 dye. **a)** Single-particle tracks in different colors. **b)** DNA-dCas9 complex zoom-in. **c)** Band-passed image from (b) for improved visualization of the protein. **d)** A repetitive (n=5) correlative recording of conductance (top) co-aligned with fluorescence intensity (bottom) measured from the location marked in (b) plotted in blue and fluorescence intensity from the background plotted in red. The conductance signal is inverted in this plot. 5 approach-retract curves were acquired at the same spot marked in (b). The yellow circle marked in (b) also signifies the approach point of the glass nanopore.

**Supplementary Figure 3.**
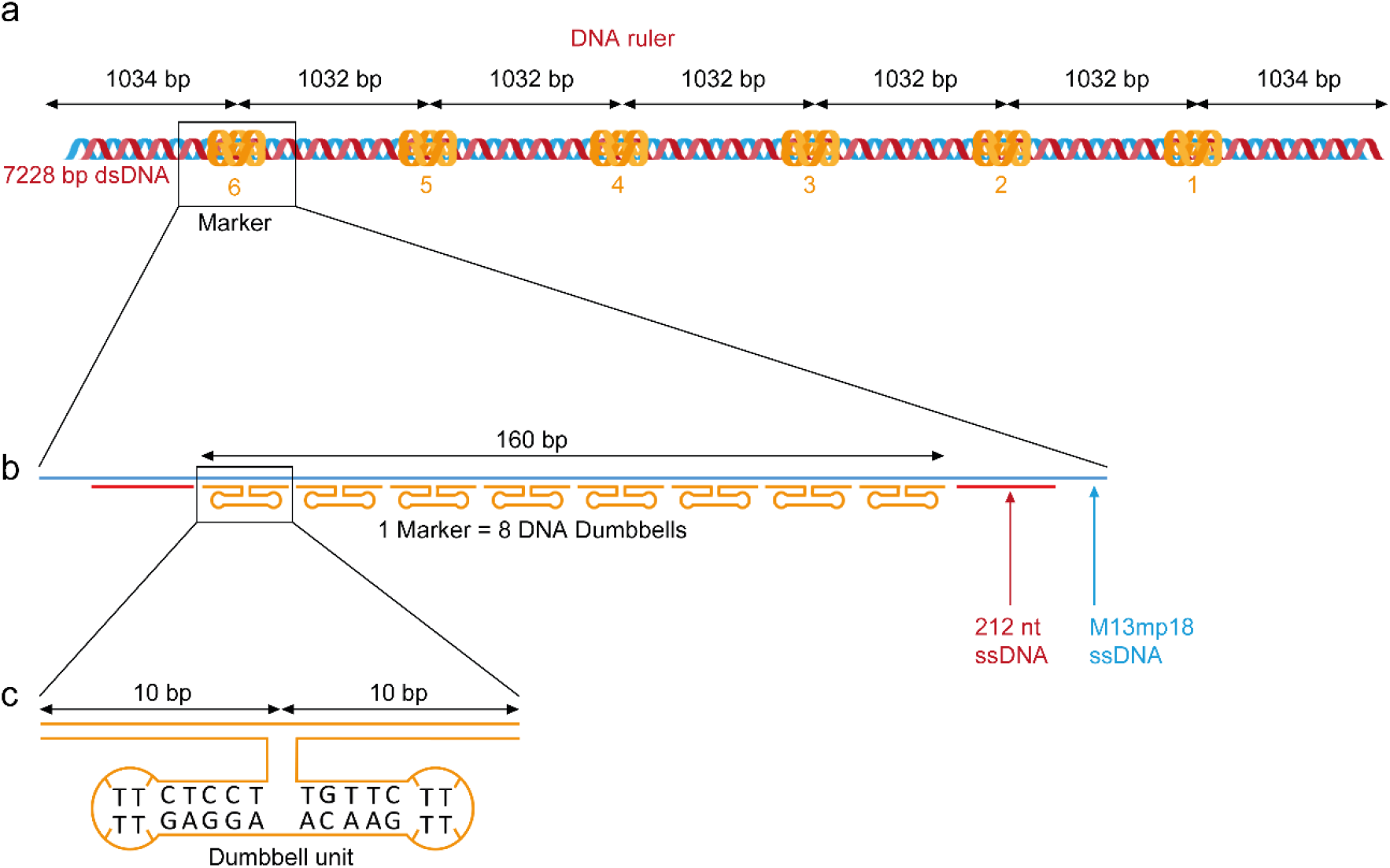
Design of the DNA ruler construct. **a)** Schematic of a DNA structure composed of 7’228 base pairs in length with 6 markers separated by equal 1032 bp intervals. **b)** Each marker contains 8 DNA dumbbell hairpin motifs (in orange), which are joined to the backbone (M13mp18 in blue). The designed dumbbells and the complementary 212 ssDNA (in red) formed the final dsDNA ruler. (see methods section for more details). **c)** Base sequence of the dumbbell hairpin motif composed of two 10 bp sections.

**Supplementary Figure 4.**
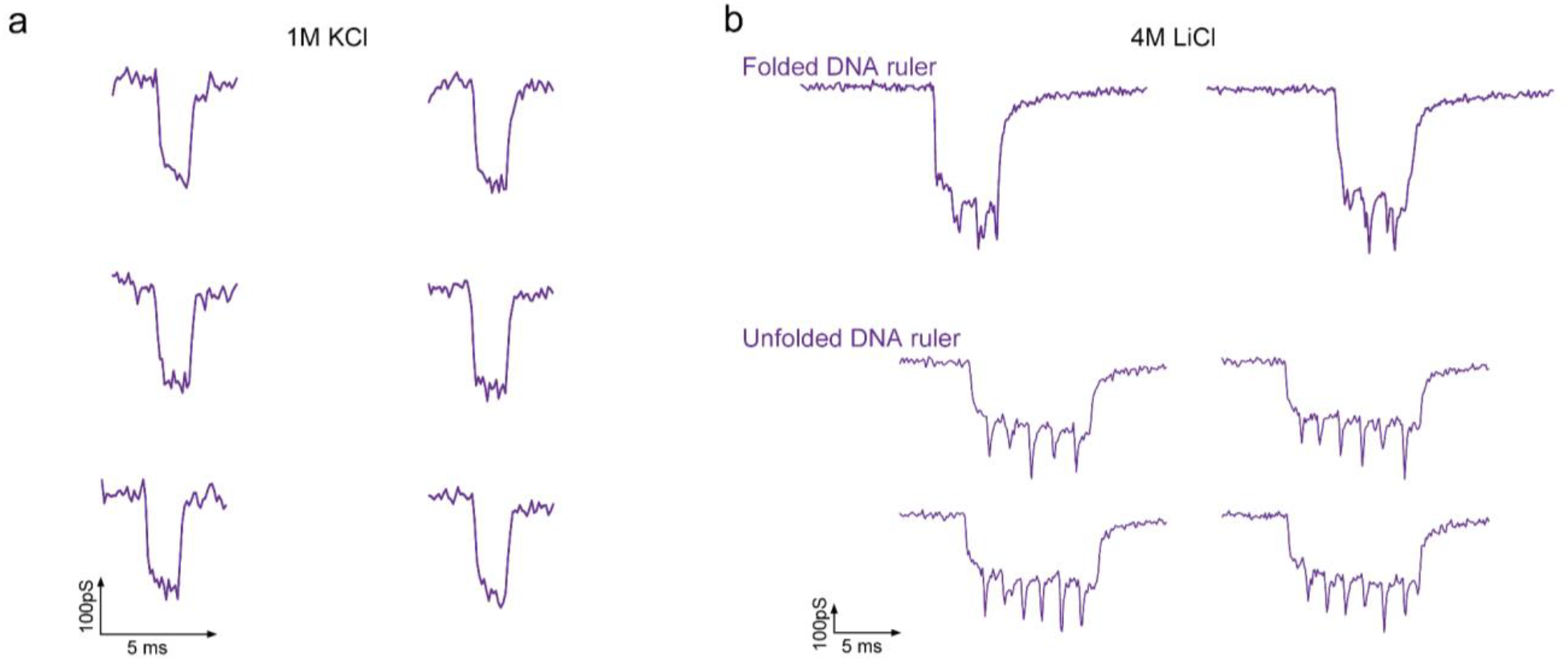
Typical free translocation curves with glass nanopores highlight the low detectability from the uncontrolled velocity. **a)** Free translocations in 1M KCl with 500 mV bias. **b)** Free translocations in 4 M LiCl. This figure highlights the challenges of free translocations in detecting topological features along DNA molecules with 600 mV bias. KCl medium has been shown to increase the SNR of the conductance signal compared with other salts (NaCl and LiCl)^13^ but the translocation speed is too high to detect the features. 4M LiCl medium is typically a good alternative to slow down translocation speed, but the detection is still limited and translocations of folded molecules are recurrent (panel b - on top).

**Supplementary Figure 5.**
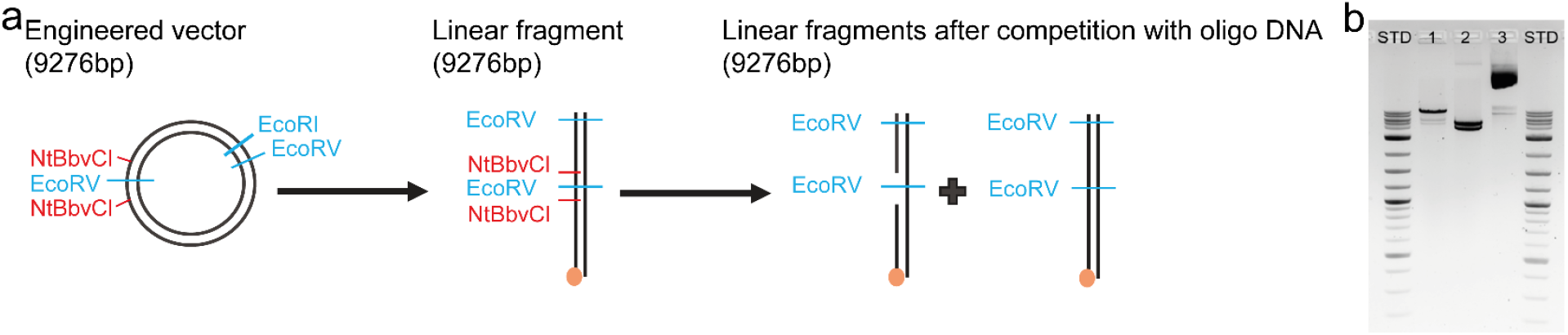
Custom-designed DNA gaps. **a)** We modified a 9 kb circular vector to introduce two Nt.BbvCI restriction sites flanking an EcoRV restriction site. First, the vector is linearized using digestion with EcoRI. Digestion with the nicking enzyme Nt.BbvCI introduces single-strand breaks in dsDNA and allows for subsequent elimination of the single-stranded DNA comprised between the nicks. The EcoRV site between Nt.BbvCI sites are used to eliminate ungapped molecules. Indeed, upon successful gap formation, EcoRV will not recognize the (now single-stranded) restriction site. In contrast, if gap formation fails and the fragment remains ungapped, digestion with EcoRV will generate two fragments of smaller sizes that can easily be eliminated. **b)** Construct characterization using electrophoresis on an agarose gel. Lane 1: Biotinylated, gapped DNA purified, after additional digestion with EcoRV. Additional digestion with EcoRV cannot cut DNA at the gap because EcoRV only hydrolyzes double-stranded DNA. Thus, the ∼9kb fragment is left intact. There might still be a minority fraction of fragments that do not contain the gap which gives rise to two smaller linear fragments (see two faint bands below the main band). Lane 2: If the engineered vector is digested with EcoRV before gap formation, it produces two linear fragments, (that appear like a single, thick band on the picture), as expected. Lane 3: Undigested engineered vector.

**Supplementary Figure 6.**
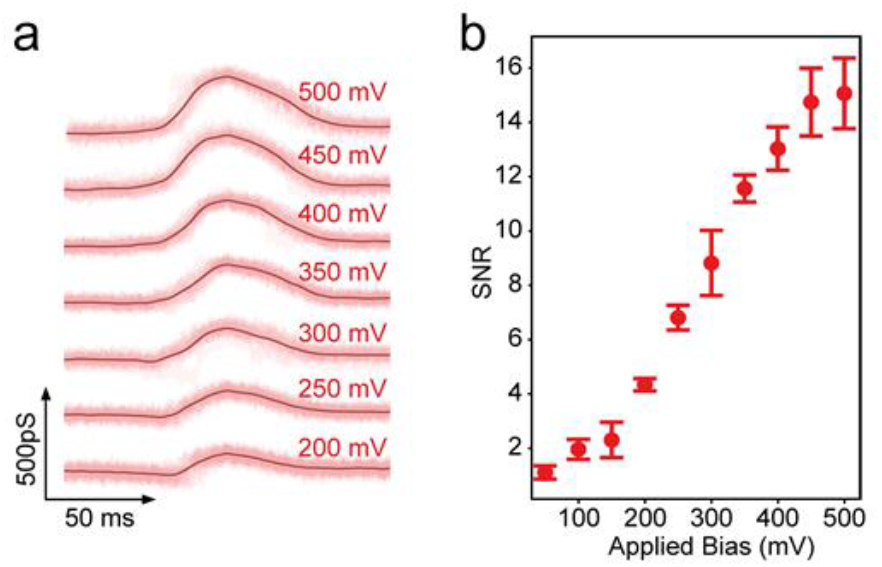
SICS translocation of the DNA gap construct containing the 80 nt gap, with a tunable bias for a constant speed of 1 μm/s. **a)** SICS curves with a 13 nm radius pipette. In dark red is the average of 10 curves for each bias. **b)** SNR of averaged translocations vs applied bias. Error bars represent the standard deviation.

**Supplementary Figure 7.**
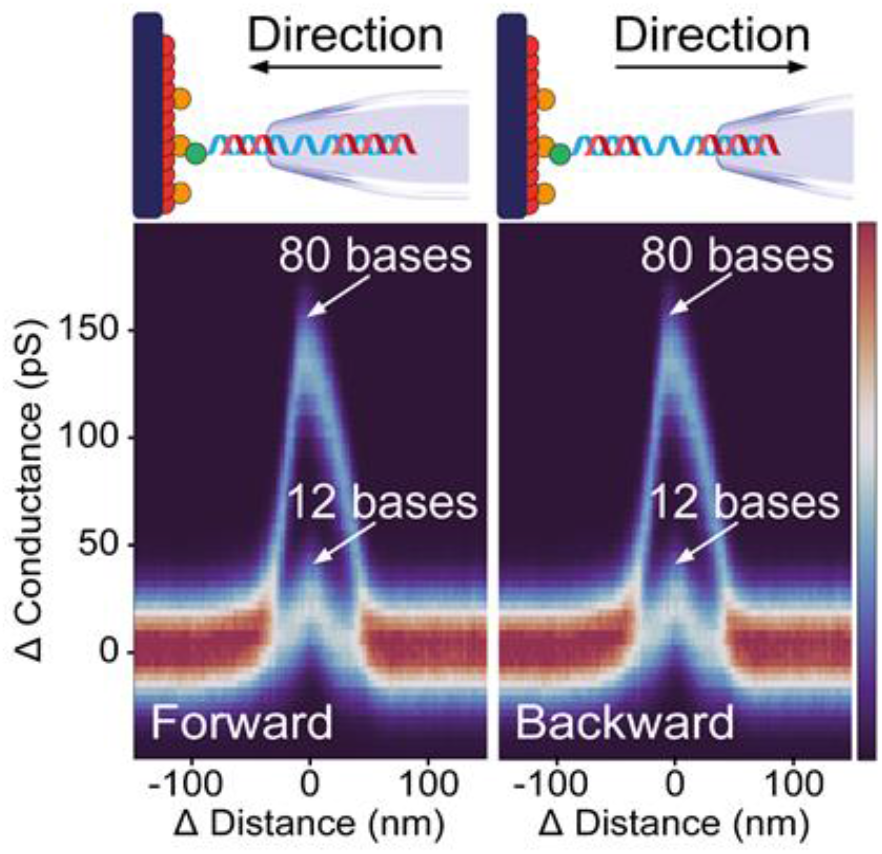
Bidirectional readings with SICS. This figure shows the color-coded probability density map (n=100 curves) of forward (left panel) and backward (right panel) controlled-translocations curves of 80 nt and 12 nt DNA gaps at 1 μm/s translocation velocity with 12 nm radius pipette. The colormap represents the normalized probability of occurrence for a conductance value at a corresponding distance. The capability to have control over the pulling directionality allowed us to observe DNA hairpins at 4 **°**C see SI Figure 10.

**Supplementary Figure 8.**
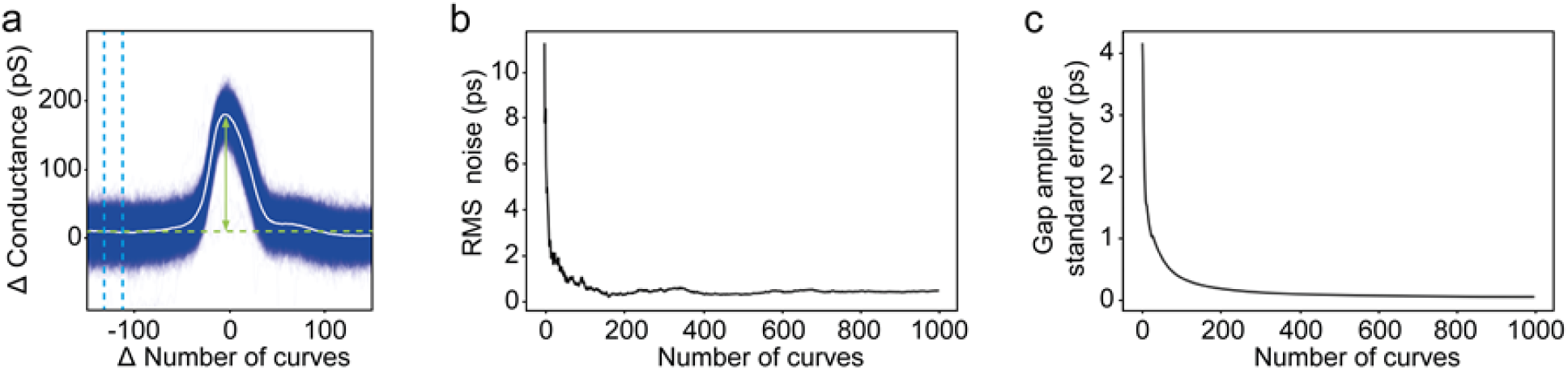
Improvement of the translocation SNR and detection of amplitude error by averaging multiple traces from the same feature on the same molecule. **a)** Average of 1000 curves on the same 80 nt gap at 1 μm/s translocation velocity with 10 nm radius pipette and 300 mV bias in 1 M KCl. **b)** RMS noise measured in 20 nm range, blue dashed lines in panel (a). **c)** Standard error of the detected gap amplitude, green arrows in panel (a).

**Supplementary Figure 9.**
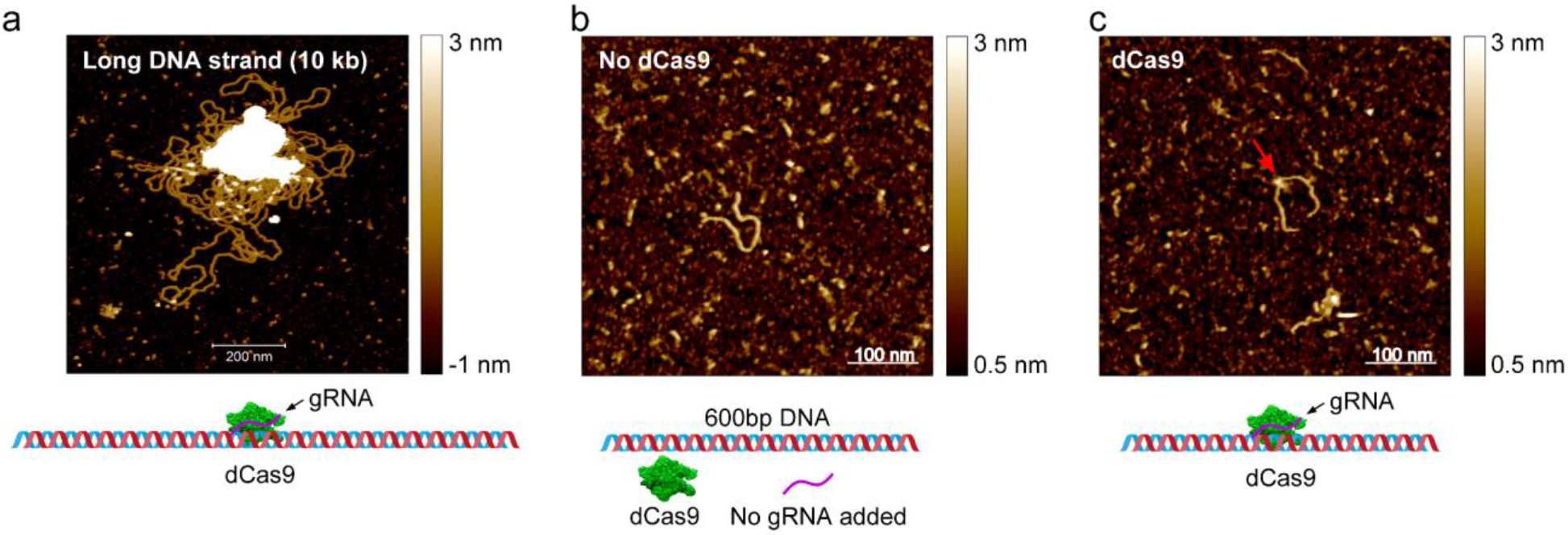
Atomic force microscopy imaging of DNA-dCas9 complex. **a)** AFM image of 10 kb long DNA incubated with dCas9 and gRNA, demonstrating that AFM analysis is not feasible on long DNA molecules. **b)** A negative control experiment with an absence of gRNA with no observed dCas9 binding. **c)** A positive control, with gRNA present in a solution with a dCas9 binding event in a field-of-view (red arrow).

**Supplementary Figure 10.**
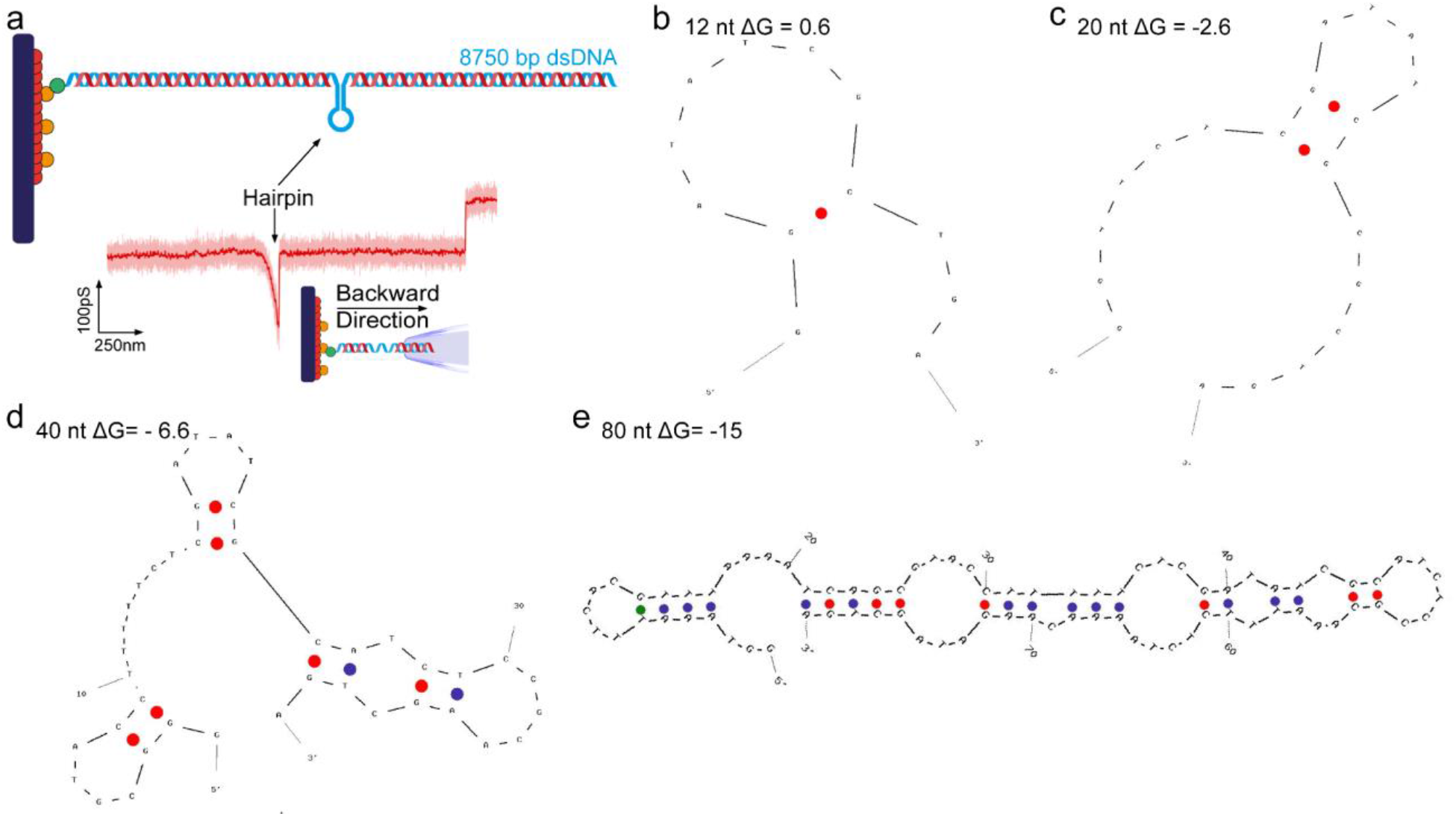
Hairpin formation in DNA gap molecules. **a)** Design of a DNA gap construct (40 nt gap) with the formation of a hairpin and a SICS conductance-distance curve in red. Prediction of secondary structures and Gibbs free energy values (ΔG) at 4**°**C for the sequences of the single-strand DNA stretch in four DNA gap constructs: 12 nucleotide gap **b)**, 20 nucleotide gap **c)**, 40 nucleotide gap **d)**, and 80 nucleotide gap **e)**. This figure highlights the formation of hairpins that were observed during SICS experiments, in samples measured right after incubation at 4**°**C. The experiments were performed with 300 mV bias in 1 M KCl. The bidirectional capability of SICS (SI Figure 7) is relevant in this application as it allows us to exploit the balance between drag and electrophoretic force. Decreasing the total force acting on the molecule in the backward direction allowed us to observe a prominent peak (a) that we assigned to the hairpin formation. In forward direction, the total force is high enough that our data are similar to the results shown in the main figures after samples have been brought to room temperature (Figure 3 and 4).

**Supplementary Figure 11.**
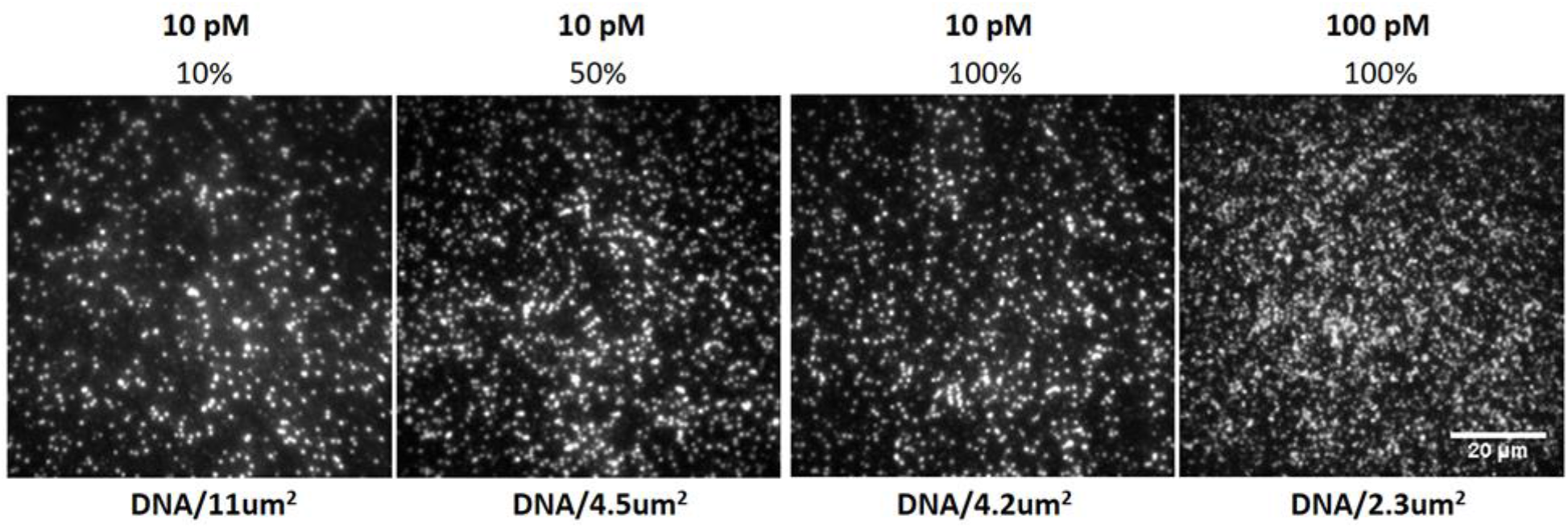
Imaging of DNA tethered on a glass coverslip with a widefield fluorescence microscope. 10 pM; 100 pM DNA solution, 5 nM; 50 nM YOYO-1 dye. Different percentage of biotinylated BSA.

**Supplementary Table 1.**
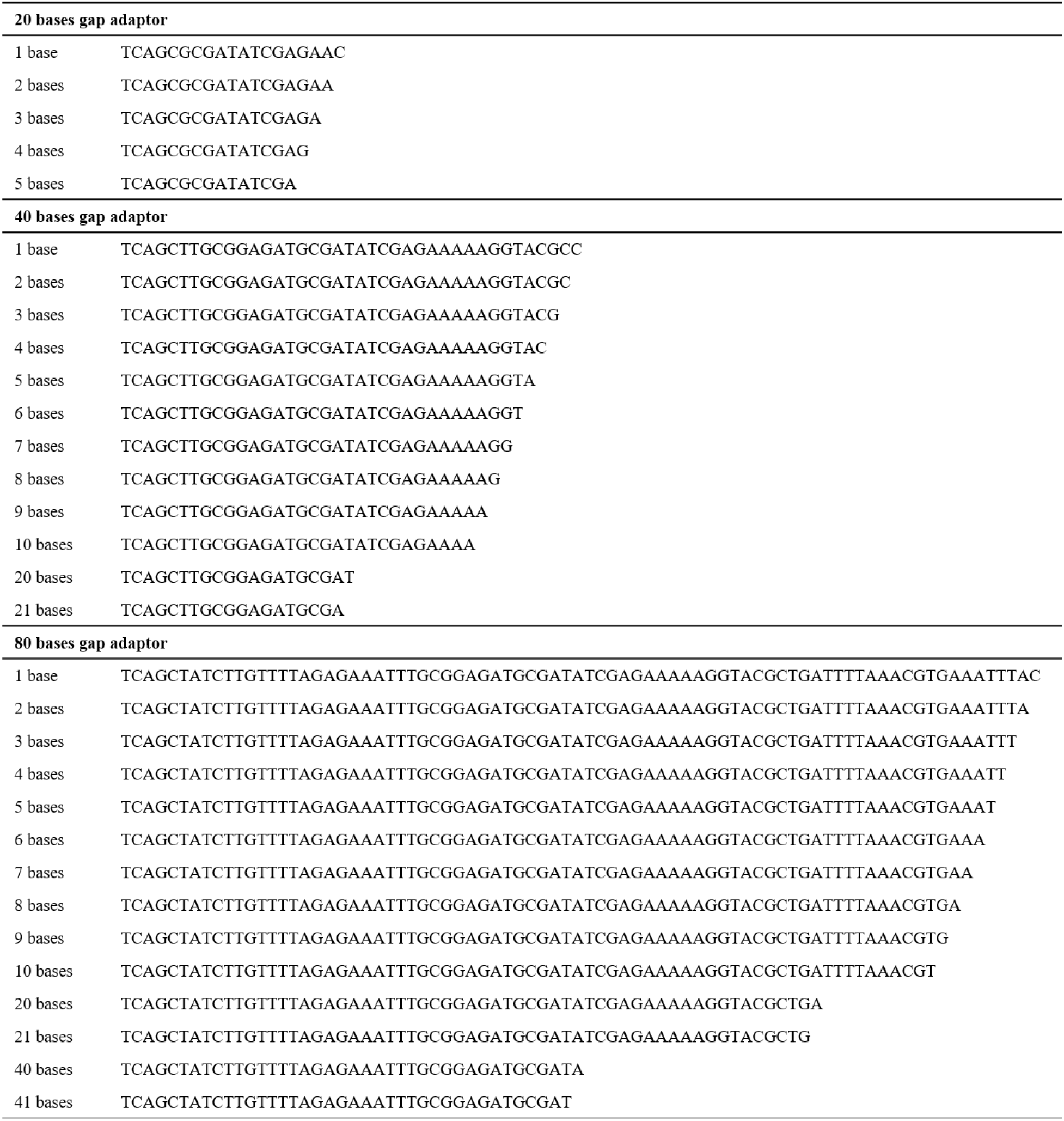
Library of gap adaptors with oligonucleotides complementary to the DNA gap constructs.

